# Malachite: A Gene Enrichment Meta-Analysis (GEM) Tool for ToppGene

**DOI:** 10.1101/511527

**Authors:** Gregory R. Gershkowitz, Zachary B. Abrams, Caitlin E. Coombes, Kevin R. Coombes

**Affiliations:** Department of Biomedical Informatics, Wexner Medical Center, The Ohio State University, Columbus, OH 43210 USA

**Keywords:** Gene enrichment analysis, Python, ToppGene

## Abstract

**Background:** Researchers commonly use online tools such as ToppGene to conduct enrichment analyses on gene expression data. This process does not easily allow multiple gene data sets to be analyzed and compared at once. ToppGene requires the user to manually enter gene symbols or other gene identifiers into a text box and to manually sift through forms with many adjustable parameters in order to obtain a downloadable text file of results. This process makes the analysis of multiple sets of genes tedious, time-consuming, and error prone. To address this problem, we developed Malachite, a Python package that enables researchers to perform gene enrichment analyses on multiple gene lists and concatenate the resulting enrichment statistics. In this way, Malachite enables meta-enrichment analyses across multiple data sets.

**Results:** To illustrate its use, we applied Malachite to three data sets from the Gene Expression Omnibus comparing gene expression in the large airways of smokers and non-smokers. Biological processes enriched in all three data sets were related to xenobiotic stimulus; molecular functions typically involved nicotinamide adenine dinucleotide phosphate (NADP) activity.

**Conclusion:** Malachite enables researchers to automate gene enrichment metaanalyses using ToppGene. Malachite also enhances ToppGene’s gene set analysis of drug-gene relationships by further filtering for FDA approved drugs.

## Background

Early in the history of gene expression profiling with microarrays, researchers recognized that it was difficult to reproduce lists of differentially expressed or predictive genes across data sets [1–4]. A small part of the problem stemmed from technological differences between the microarray platforms used to perform the assays. However, other challenges were more fundamental and remain relevant today after replacing microarrays with RNA sequencing. Individual false positive genes, tumor heterogeneity, small sample sizes, and highly correlated genes all contribute to instability in lists of selected genes [5–7].

Efforts to deal with these challenges evolved in two directions. Some researchers hypothesized that correlation could be explained because genes were active in the same pathway or as part of the same biological process, and incorporated the tools of gene set enrichment analysis [8–10] into their attempts to understand differential expression or develop predictive models [11–13]. Other researchers focused on increasing the effective sample size by applying meta-analysis to multiple related data sets [14–16]. More recently, several groups have combined meta-analysis with gene set enrichment analysis. As with traditional single-measure meta-analyses, these include methods that require all of the original data [17], methods that work with summary statistics [18,19], and methods that work with p-values [20–22].

The need for biologists to perform their own meta-analyses of gene set enrichment has increased over time. For example, the Gene Expression Omnibus (GEO), which at this writing contains more than 100,000 data sets, introduced GEO2R in 2013 [23]. GEO2R allows users to perform differential expression analyses online, easily retrieving lists of genes with associated t-statistics and adjusted p-values. However, comparing the results of experiments can be difficult due to batch effects or differences between experimental platforms. Manually entering each resulting gene list into an online tool for gene set enrichment analysis can be both tedious and error-prone if researchers want to use consistent non-default parameters. There are many online tools to perform gene set enrichment analysis [24]. However, to the knowledge of the authors, there is no tool that enables users to select which category (for example, drugs, gene ontology (GO) terms, or cytogenetic bands) of enrichment analysis to perform uniformly over a group of distinct data sets, and concatenate the results for meta-analysis. This additional step would enable researchers to target their enrichment analyses to better suit their experimental needs. It would also streamline the process of performing enrichment analysis, since multiple data sets could be studied at the same time.

In this note, we describe Malachite, a Python package that allows users to easily collect gene lists from multiple data sets, perform gene set enrichment analysis using a consistent set of parameters, and combine the resulting lists of gene set enrichment results across multiple enrichment categories (Figure 1). Malachite relies on ToppGene (https://ToppGene.cchmc.org/) for the gene set enrichment analysis of individual gene lists [8]. ToppFun, a component of ToppGene, performs functional enrichment analyses based on underlying transcriptome, ontology, phenotype, proteome, and pharmacome annotations. Using Malachite, researchers can select multiple lists of significant gene data, set the enrichment analysis parameters, and run each list through ToppGene. Malachite then concatenates the results in an automated fashion, allowing for further meta-analysis. Additional filtering by FDA drugs approved to treat cancer is performed if the user selected “drugs” as one of their enrichment categories. Thus, Malachite enhances ToppGene’s gene set analysis of drug-gene relationships by allowing users to focus on FDA-approved cancer drugs.

**Figure 1:**
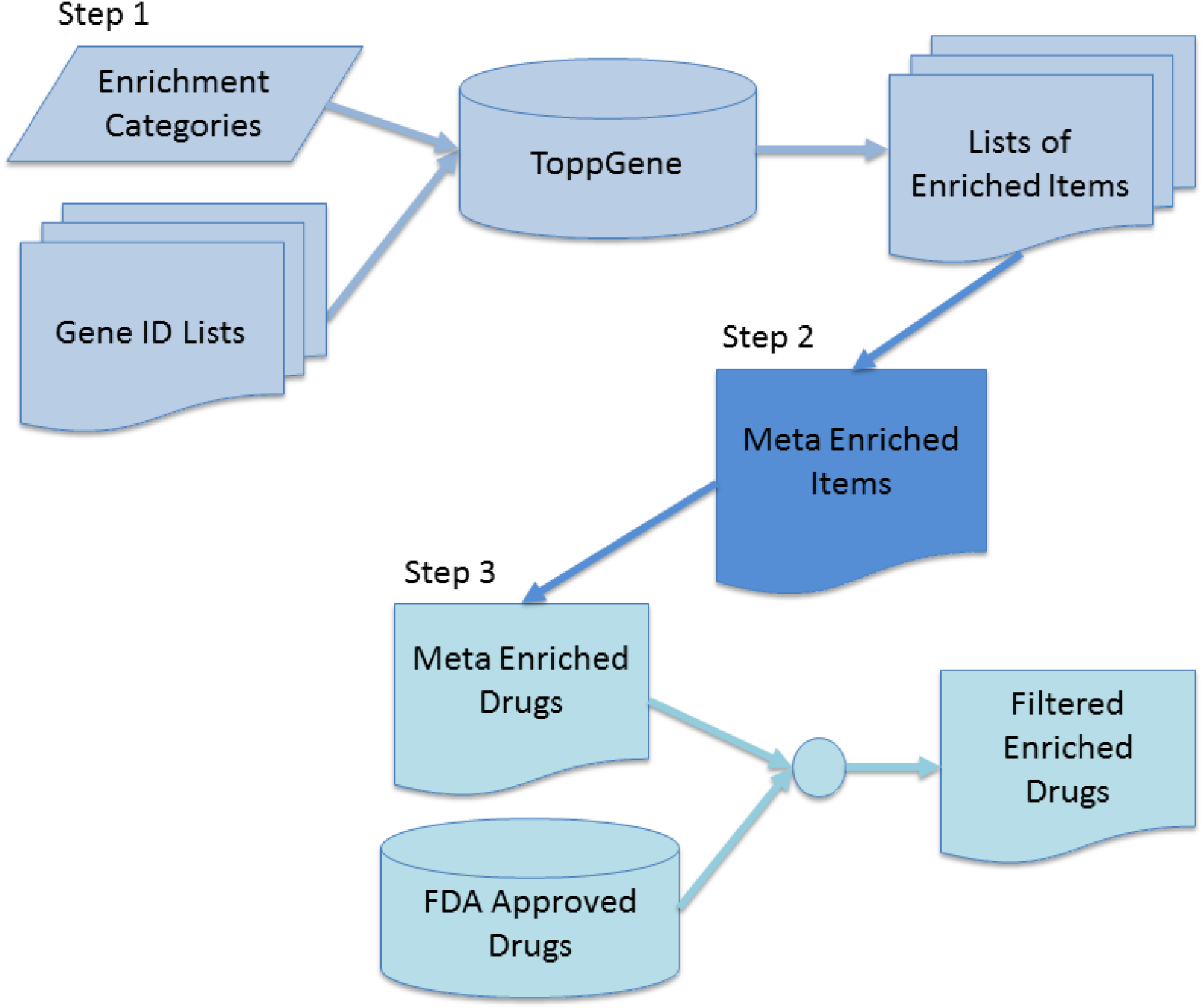
Malachite workflow. Step1: The user specifies the enrichment categories of interest and the type of gene identifier used, along with other parameters. Malachite combines these specifications with the gene lists and passes each list to ToppGene. The set of enriched items from each list are saved. Step 2: Malachite automatically performs meta-analysis, recording the number of lists in which each term is found to be significant. Step 3: If one of the chosen categories is “Drugs”, then Malachite automatically filters the list of enriched drugs against the database of FDA-approved drugs.

## Implementation

Malachite is a Python package that is designed to automate and correlate multiple independent ToppGene enrichment analyses. First, the program invokes ToppFun, using Mechanize, a Python module used for web crawling. Malachite selects the type of gene identifier (Entrez ID or HGNC symbol) specified by the user and submits the form. Second, Malachite saves the unique session ID for the user using Beautiful Soup, a Python library for retrieving data from a web site. Third, Malachite, via Mechanize, opens the URL with the session ID and specifies which database categories (for example, “Disease” and/or “Drug”) the user would like to analyze. Fourth, Malachite causes ToppGene to run each analysis and downloads the resulting text files of results. Finally, Malachite processes the multiple tab-delineated results files by database category and combines the results.

In addition, Malachite filters gene set enrichment analyses using drug databases. By comparing drugs to FDA lists (https://www.cancer.gov/about-cancer/treatment/drugs), Malachite can identify FDA approved cancer therapies. This important step helps filter down the list of potential drug targets, especially for applications such as drug repositioning [25]. This FDA filtering step aids in translating potential meta-analysis findings into actionable results.

## Results

To illustrate Malachite, we applied it to three Gene Expression Omnibus (GEO) datasets (GSE5056, GSE5057 and GSE5059), each of which used a different generation of Affymetrix microarrays to test expression differences in epithelial samples obtained by bronchoscopy from the large airways of smokers and nonsmokers [26]. Gene lists were obtained using the GEO2R facility at the GEO web site; these gene lists, along with instructions for how to reproduce them, are contained in Additional File 1. The data sets were processed by Malachite for drugs, diseases, and gene ontology terms. Although 86 drugs or small molecules were enriched in all three data sets, none of them were FDA-approved for the treatment of cancer. No diseases were enriched in all three data sets. The GO terms for “Biological Process” and “Molecular Function” are listed in Table 1. The only Biological Process terms that were found in all three datasets were related to xenobiotic stimulus, representing the foreign nature of chemical damage from smoking. The Molecular Function terms show enrichment for nicotinamide adenine dinucleotide phosphate (NADP) activity. This results shows that Malachite can perform enrichment metaanalyses across datasets and recover the underlying biological content.

**Table 1:**
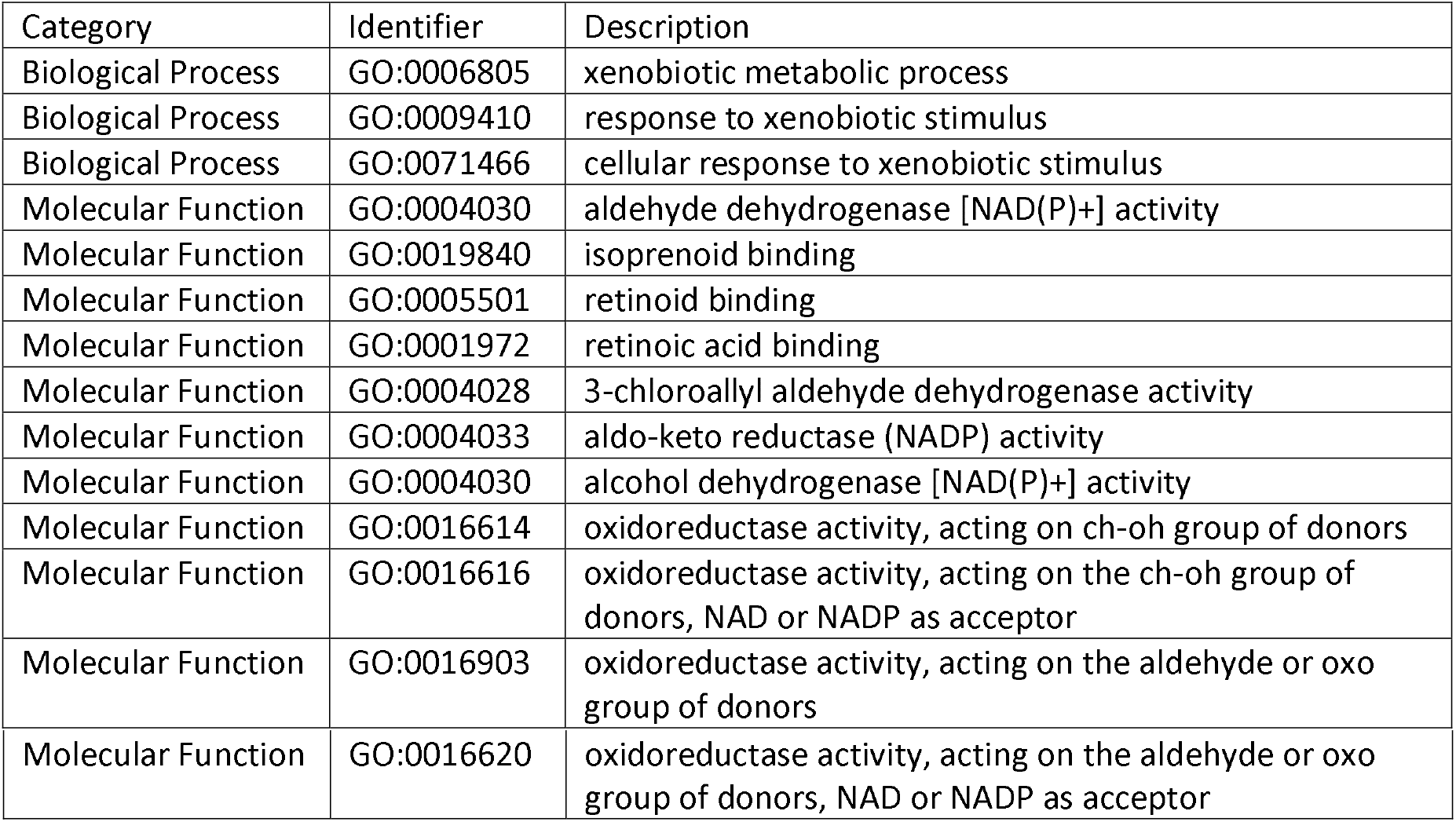
Gene Ontology terms enriched in all three data sets comparing smokers to non-smokers.

## Conclusion

Currently, if researchers want to perform gene enrichment analysis on multiple gene lists using ToppGene, they must go through the manual process of selecting the many user specifications the site provides, as well as a painstaking process of combining the enrichment data for all sets of results. We developed Malachite to allow ToppGene users to input multiple lists of genes, set the enrichment analysis parameters, and run these sets through ToppGene to obtain results, which are stored in individual text files. Malachite then concatenates the results in an automated fashion, allowing for further meta-data analyses, with special attention paid to post-processing lists of associated drugs.

## Availability and Requirements

Project name: Malachite

Project homepage: at https://pypi.org/project/malachite

Operating system(s): Platform independent

Programming language: Python

Other requirements: NA

License: GNU

## List of abbreviations

HGNC: : HUGO Gene Nomenclature Committee
GEO: : Gene Expression Omnibus
GO: : Gene ontology

## Declaration

### Ethics approval and consent to participants

Not applicable

### Consent for publication

Not applicable

### Availability of data and material

All data generated or analyzed during this study are included in this article and its additional files.

### Author’s contributions

Gregory R. Gershkowitz: Lead software developer and wrote the Python code for the malachite package and helped write the paper.

Zachary B. Abrams: Helped design and develop software for the malachite package and was the lead in writing the paper.

Caitlin E. Coombes: Helped edit and write the paper.

Kevin R. Coombes: Oversaw all aspects of the project e.g. Software design, software development, paper writing and editing.

## Acknowledgments

We thank the Summer Internship Program at the Ohio State University Department of Biomedical Informatics for their support.

## Additional files

Additional File 1: Supplementary material. This file contains detailed instructions on how to use Malachite (ZIP 2876 KB), structured as a mini-website. This compressed file includes:

- Index.html (source readme.md): an overview of the process
- UseGEO2R.html (source: UseGEO2R.md): instructions for using the UseGEo2R tool at the Gene Expression Omnibus.
- PrepForMalachite.html (source: PrepForMalachite.ipynb): an example Jupyter notebook showing how to prepare input files.
- RunMalachite.html (source: RunMalachite.ipynb): an example Jupyter notebook showing how to run Malachite.
- Three input files obtained by using GEO2R (GSE5056_Smoking_Malachite.csv, GSE5057_Smoking_Malachite.csv, and GSE5059_Smoking_Malachite.csv)
- Two auxiliary files to make the html files from their sources (Makefile.Win, dashed.css).

## References

1. Ein-Dor, L., et al., Outcome signature genes in breast cancer: is there a unique set? Bioinformatics, 2005. 21(2): p. 171–8.

2. Lossos, I.S., et al., Prediction of survival in diffuse large-B-cell lymphoma based on the expression of six genes. N Engl J Med, 2004. 350(18): p. 1828–37.

3. Miklos, G.L. and R. Maleszka, Microarray reality checks in the context of a complex disease. Nat Biotechnol, 2004. 22(5): p. 615–21.

4. Ntzani, E.E. and J.P. Ioannidis, Predictive ability of DNA microarrays for cancer outcomes and correlates: an empirical assessment. Lancet, 2003. 362(9394): p. 1439–44.

5. Ma, S., et al., Integrative analysis of multiple cancer prognosis studies with gene expression measurements. Stat Med, 2011. 30(28): p. 3361–71.

6. Michiels, S., S. Koscielny, and C. Hill, Interpretation of microarray data in cancer. Br J Cancer, 2007. 96(8): p. 1155–8.

7. Xu, J.Z. and C.W. Wong, Hunting for robust gene signature from cancer profiling data: sources of variability, different interpretations, and recent methodological developments. Cancer Lett, 2010. 296(1): p. 9–16.

8. Chen, J., et al., ToppGene Suite for gene list enrichment analysis and candidate gene prioritization. Nucleic Acids Res, 2009. 37(Web Server issue): p. W305–11.

9. Huang da, W., B.T. Sherman, and R.A. Lempicki, Systematic and integrative analysis of large gene lists using DAVID bioinformatics resources. Nat Protoc, 2009. 4(1): p. 44–57.

10. Subramanian, A., et al., Gene set enrichment analysis: a knowledge-based approach for interpreting genome-wide expression profiles. Proc Natl Acad Sci U S A, 2005. 102(43): p. 15545–50.

11. Abraham, G., et al., Prediction of breast cancer prognosis using gene set statistics provides signature stability and biological context. BMC Bioinformatics, 2010. 11: p. 277.

12. Manoli, T., et al., Group testing for pathway analysis improves comparability of different microarray datasets. Bioinformatics, 2006. 22(20): p. 2500–6.

13. Zhang, M., et al., Evaluating reproducibility of differential expression discoveries in microarray studies by considering correlated molecular changes. Bioinformatics, 2009. 25(13): p. 1662–8.

14. Park, W.D. and M.D. Stegall, A meta-analysis of kidney microarray datasets: investigation of cytokine gene detection and correlation with rt-PCR and detection thresholds. BMC Genomics, 2007. 8: p. 88.

15. Wang, J., et al., Differences in gene expression between B-cell chronic lymphocytic leukemia and normal B cells: a meta-analysis of three microarray studies. Bioinformatics, 2004. 20(17): p. 3166–78.

16. Yang, X. and X. Sun, Meta-analysis of several gene lists for distinct types of cancer: a simple way to reveal common prognostic markers. BMC Bioinformatics, 2007. 8: p. 118.

17. Chen, M., et al., A powerful Bayesian meta-analysis method to integrate multiple gene set enrichment studies. Bioinformatics, 2013. 29(7): p. 862–9.

18. Shen, K. and G.C. Tseng, Meta-analysis for pathway enrichment analysis when combining multiple genomic studies. Bioinformatics, 2010. 26(10): p. 1316–23.

19. Zhang, H., et al., A Powerful Procedure for Pathway-Based Meta-analysis Using Summary Statistics Identifies 43 Pathways Associated with Type II Diabetes in European Populations. PLoS Genet, 2016. 12(6): p. e1006122.

20. Kaever, A., et al., Meta-analysis of pathway enrichment: combining independent and dependent omics data sets. PLoS One, 2014. 9(2): p. e89297.

21. Langfelder, P., P.S. Mischel, and S. Horvath, When is hub gene selection better than standard meta-analysis? PLoS One, 2013. 8(4): p. e61505.

22. Nguyen, T., et al., A novel bi-level meta-analysis approach: applied to biological pathway analysis. Bioinformatics, 2016. 32(3): p. 409–16.

23. Barrett, T., et al., NCBI GEO: archive for functional genomics data sets--update. Nucleic Acids Res, 2013. 41(Database issue): p. D991–5.

24. Huang, D.W., B.T. Sherman, and R.A. Lempicki, Bioinformatics enrichment tools: paths toward the comprehensive functional analysis of large gene lists. Nucleic acids research, 2008. 37(1): p. 1–13.

25. Ashburn, T.T. and K.B. Thor, Drug repositioning: identifying and developing new uses for existing drugs. Nature reviews Drug discovery, 2004. 3(8): p. 673.

26. Carolan, B.J., et al., Up-regulation of expression of the ubiquitin carboxyl-terminal hydrolase L1 gene in human airway epithelium of cigarette smokers. Cancer Res, 2006. 66(22): p. 10729–40.

