## Additional File 1 for "Malachite: A Gene Enrichment Meta-Analysis (GEM) Tool for ToppGene": index.html

Malachite Supplement


### Malachite Supplement

#### Greg Gershkowitz and Kevin R. Coombes

##### 4 April 2019

### Overview

This folder contains supplementary material for our paper describing the Malachite Python package for performing Gene Enrichment Meta-analysis (GEM) starting with multiple lists of genes.

#### Input Files

There are three example input files:

1. GSE5056\_Smoking\_Malachite.csv
2. GSE5057\_Smoking\_Malachite.csv
3. GSE5059\_Smoking\_Malachite.csv

These input files were produced using the Gene Expression Omnibus (GEO) web site by running the GEO2R tool to compare microarray data from smokers and non-smokers under various conditions. (For more details,see UseGEO2R.md).

#### Preparation

To prepare the data for input into Malachite, it must be filtered and collected into a single Excel spreadsheet, where each column contains a separate list of genes. An example of how to do that is given in a Jupyter notebook, which can be viewed in HTML format. The resulting Excel spreadsheet is called `forMalachite.xls`.

#### Running Malachite

You can then run Malachite from a command line. To illustrate the comand arguments, we have provided another Jupyter notebnook, which can also be viewed in HTML format.
