## Additional File 1 for "Malachite: A Gene Enrichment Meta-Analysis (GEM) Tool for ToppGene": PrepForMalachite.html


### Preparing to Run Malachite¶

#### Getting Started¶

**Note:** In order to write to Excel files, you need to install the module 'openpyxl', which at least for me was not automatically installed when I used 'pip' to install 'pandas'.

In [1]:

```
import pandas
import openpyxl
import glob
import os
```

This script assumes that we want to use all of the CSV files in the current working directory.

In [2]:

```
csvfiles = glob.glob("*.csv")
csvfiles
```

Out[2]:

```
['GSE5056_Smoking_Malachite.csv',
 'GSE5057_Smoking_ Malachite.csv',
 'GSE5059_Smoking_Malachite.csv']
```

#### Illustrating the procedure with one file¶

The basic idea is to use the 'pandas' module to both read CSV files and write Excel files. We are going to illustr5ate this process with the first CSV file.

In [3]:

```
f = csvfiles[1]
nc = f.find("_") # find the first underscore
gse = f[:nc]     # The GSE identifier is the part before the underscore
gse
```

Out[3]:

```
'GSE5057'
```

Now we can read the file.

In [4]:

```
df = pandas.read_csv(f)
df.shape
```

Out[4]:

```
(22283, 8)
```

As expected, we got the full spreadsheet, with more than 22,000 gene-rows and 8 columns. Next, we take a look at the column names.

In [5]:

```
df.dtypes
```

Out[5]:

```
AffyID          object
adj.P.Val      float64
P.Value        float64
t              float64
B              float64
logFC          float64
Gene.symbol     object
Gene.title      object
dtype: object
```

We want to select rows based on the adjusted p-value.

In [6]:

```
df2 = df.loc[df['adj.P.Val'] < 0.10]
df2.shape
```

Out[6]:

```
(27, 8)
```

Only 27 rows meet our selection criterion.

Next, we can write the selected rows to an Excel spreadsheet.

In [7]:

```
df2.to_excel("test.xlsx", sheet_name = gse, index = False)
```

#### Repeat for all CSV files¶

Here, we perform the same read-filter-write operation on all CSV files in the directory. This process is somewhat more complicated since we want to write all the information to the same spreadsheet, with each source file going to a separate worksheet.

In [8]:

```
fname = "forMalachite.xlsx"
try:
    os.remove(fname)
except OSError:
    pass
writer = pandas.ExcelWriter(fname, engine="openpyxl")
if os.path.exists(fname):
    book = openpyxl.load_workbook(fname)
    sheet_names = book.sheet_names()
    print('Sheet Names', sheet_names)
    writer.book = book
writer.path
```

Out[8]:

```
'forMalachite.xlsx'
```

In [9]:

```
lof = [] # list of file contents
lon = [] # list of names
for f in csvfiles:
    nc = f.find("_")
    gse = f[:nc]
    lon.append(gse)
    df = pandas.read_csv(f)
    df.dtypes
    df2 = df.loc[df['adj.P.Val'] < 0.10]
    lof.append(df2)
    df2.shape
    df2.to_excel(writer, sheet_name = gse)
    writer.save()
lon
```

Out[9]:

```
['GSE5056', 'GSE5057', 'GSE5059']
```

Note that we have not yet *closed* the workbook. That's because we want to put together one more worksheet (at rthe beginning) that combines information from all of the pieces we have looked at so far.

In [10]:

```
result = writer.book.create_sheet("0.1Cutoff", 0) # the '0' puts this sheet in front of all others
rowfix = 1
columnfix = 1
for name in lon:
    result.cell(row=rowfix, column=columnfix).value = str(name)
    columnfix += 1
writer.sheets # why doesn't this show all four sheets? They end up in the actuaol Excel file...
```

Out[10]:

```
{'GSE5056': <Worksheet "GSE5056">,
 'GSE5057': <Worksheet "GSE5057">,
 'GSE5059': <Worksheet "GSE5059">}
```

In [11]:

```
columnfix = 1
for file in lof:
    print(file.shape)
    num_rows = file.shape[0]
    num_cols = file.shape[1]
    rowfix = 2
    for i in range(1, num_rows):
        result.cell(row=rowfix, column=columnfix).value = file.at[i,'Gene.symbol']
        rowfix += 1
    columnfix += 1
```

```
(177, 8)
(27, 8)
(10, 8)
```

In [12]:

```
writer.close()
```
