## Additional File 1 for "Malachite: A Gene Enrichment Meta-Analysis (GEM) Tool for ToppGene": Readme.md

---
author: Greg Gershkowitz and Kevin R. Coombes
date: 4 April 2019
title: Malachite Supplement
---

# Overview
This folder contains supplementary material for our paper describing
the Malachite Python package for performing Gene Enrichment
Meta-analysis (GEM) starting with multiple lists of genes.

## Input Files
There are three example input files:

1. [GSE5056_Smoking_Malachite.csv](./GSE5056_Smoking_Malachite.csv)
2. [GSE5057_Smoking_Malachite.csv](./GSE5057_Smoking_Malachite.csv)
3. [GSE5059_Smoking_Malachite.csv](./GSE5059_Smoking_Malachite.csv)

These input files were produced using the Gene Expression Omnibus (GEO)
web site by running the GEO2R tool to compare microarray data from
smokers and non-smokers under various conditions. (For more details,see
[UseGEO2R.md](./UseGEO2R.md)).

## Preparation
To prepare the data for input into Malachite, it must be filtered and
collected into a single Excel spreadsheet, where each column contains
a separate list of genes. An example of how to do that is given in a
[Jupyter notebook](./PrepForMalachite.ipynb), which can be viewed in
[HTML format](./PrepForMalachite.html).
The resulting Excel spreadsheet is called `forMalachite.xls`.

## Running Malachite
You can then run Malachite from a command line. To illustrate the
comand arguments, we have provided another [Jupyter
notebnook](./RunMalachite.ipynb), which can also be viewed in [HTML
format](./RunMalachite.html).
