## Additional File 1 for "Malachite: A Gene Enrichment Meta-Analysis (GEM) Tool for ToppGene": RunMalachite.ipynb

```
# %% [markdown]
# Running Malachite
## Input File

The main input to the Malachite program is an Excel spreadsheet whose first worksheet contains lists of genes in each coilumn. The current implementation requires that you supply an absolute path to this file, not a relative path. Since the file we want to use was produced in the current directory, we can figure out where that is without having to type it ourselves.

# %% [code]
import os 
thisDir = os.getcwd()
thisDir

# %% [markdown]
Now we can append the (not terribly original) file name to the current path to describe the input file.

# %% [code]
inputFile = os.path.join(thisDir, "forMalachite.xlsx")
inputFile

# %% [markdown]
## Getting Ready for Output
Malachite produces output results for each individual gene list (column in the input), which are obtained by calling ToppGene. It then makes a second pass to integrate this information, and stores those combined results somewhere. Each set of output data needs a directory where it will be stored. The next commands create these directories if they do not already exist.

# %% [code]
if not os.path.exists("Iout"):
    os.mkdir("Iout")
if not os.path.exists("Cout"):
    os.mkdir("Cout")

# %% [markdown]
## Setting Other Options
You also have to tell Malachiote what kinds of identifiers are used to indicate the input genes. Our file uses the standard gene symbols defined by the Human Gene Nomenclature Committee. (The spaces are included to make it easier to build the command string later on.

# %% [code]
geneIdentifier = " HGNC "

# %% [markdown]
You also have to specify what output categories you want to retrieve from ToppGene. We are going to collect data on associated diseases and drugs.

# %% [code]
categories = " ['Disease','Drug','GeneOntologyBiologicalProcess','GeneOntologyMolecularFunction','Pathway'] "

# %% [markdown]
## Running Malachite
Now we are going to put together the command line. You could, of course, just type this command directly in a command window, but we are going to call it from within this (Jupyter) Python notebook as a "system call".

# %% [code]
command = "malachite " + inputFile + geneIdentifier + categories + " Iout/ " + " Cout/"
command

# %% [markdown]
And we can finally run the program.

# %% [code]
os.system(command)

# %% [markdown]
A return value of 0 means that things worked without an (obvious?) error. We can list the files in the directories to get an idea of what was produced.

# %% [code]
os.listdir(os.path.join(thisDir, "Iout"))

# %% [code]
os.listdir(os.path.join(thisDir, "Cout"))
```
