## Additional File 1 for "Malachite: A Gene Enrichment Meta-Analysis (GEM) Tool for ToppGene": UseGEO2R.html

Using GEO2R


### Using GEO2R

#### Greg Gershkowitz and Kevin R. Coombes

##### 4 April 2019

### Overview

This file contains instructions for running the supplemental data experiment on smokers versus non-smokers for GSE5056, GSE5057 and GSE5059.

1. Go to the NCBI Gene Expression Omnibus page for one of the three datasets (GSE5056, GSE5057, and GSE5059).
2. Click the button labeled "Analyze using GEO2R" at the bottom of the page.
3. Click on the link labeled "Define groups" and create two groups: smokers and non-smokers.
4. Select all samples from the drop-down list of samples that are from the same group.
5. With those samples selected, click on the appropriate group from the "Define groups" tab. Repeat for both groups.
6. Click the "Save all results" link; this action will open all of the results in a new browser tab.
7. Right click on this page and save it as a text file. This action will produce a file in tab-separated-values text format, with every cell (including the numeric ones) in quotation marks. We temporarily saved the reuslts of the three analyses as
   - GEO5056\_Smoking\_Malachite.txt,
   - GEO5057\_Smoking\_Malachite.txt, and
   - GEO5059\_Smoking\_Malachite.txt.
8. By opening one of these files in Excel you can save them in comma-separated-values (CSV) format. A key difference is that Excel automatically recognizes the numeric columns and gets rid of the quotation marks when you do this.

**These files are provided in our supplemental data as GEO5056\_Smoking\_Malachite.csv, GEO5057\_Smoking\_Malachite.csv and GEO5059\_Smoking\_Malachite.csv.**

**Warning:** Trying to open the CSV files (which Excel created) in Excel will genearate a waring message. Excel will compain that the files are not in the expected format, and wlll then complain that it thinks they are "SYLK" files. Excel does this because that it thinks that any file whose first column is named "ID" is actually a "SYLK" file (https://en.wikipedia.org/wiki/SYmbolic\_LinK\_(SYLK)). With great difficulty, we will refrain from commenting further on programmers who create a file that they themselves cannot then correctly recognize.
